# Impacted Spike Frequency Adaptation Associated with Reduction of KCNQ2/3 Promotes Seizure Activity in Temporal Lobe Epilepsy

**DOI:** 10.1101/2020.09.25.313254

**Authors:** Rongrong Li, Shicheng Jiang, Shuo Tan, Bei Liu, Yang Liu, Lei Jiang, Hong Ni, Qiyi Wang, Shidi Zhao, Hao Qian, Rongjing Ge

## Abstract

Although numerous epilepsy-related genes have been identified by unbiased genome-wide screening based on samples from both animal models and patients, the druggable targets for temporal lobe epilepsy (TLE) are still limited. Meanwhile, a large number of candidate genes that might promote or inhibit seizure activities are waiting for further validation. In this study, we first analyzed two public databases and determined the significant down-regulations of two M-type potassium channel genes (KCNQ2/3) expressions in hippocampus samples from TLE patients. Then we reproduced the similar pathological changes in the pilocarpine mouse model of TLE and further detected the decrease of spike frequency adaptation driven by impacted M-currents on dentate gyrus granule neurons. Finally, we employed a small-scale simulation of dentate gyrus network to investigate potential functional consequences of disrupted neuronal excitability. We demonstrated that the impacted spike frequency adaptation of granule cells facilitated the epileptiform activity among the entire network, including prolonged seizure duration and reduced interictal intervals. Our results identify a new mechanism contributing to ictogenesis in TLE and suggest a novel target for the anti-epileptic drug discovery.

## INTRODUCTION

Epilepsy is a group of devastating neuronal disorders characterized as recurrent seizure activities[1]. The functional disruptions of dentate gyrus (DG) are believed to be crucial in the generation of epileptic seizures in the patients with temporal lobe epilepsy[2–4]. Several key dysfunctions of dentate granule cells engaging in the hippocampal sclerosis enhance the hyperexcitability of DG and entire hippocampus network, involving mossy fiber back-sprouting and reduction of interneurons[5, 6]. Besides those pathological alternations of neuronal network, the instinct electrophysiological properties of granule cells are also documented as a part of mechanisms promoting ictogenesis[7–9]. Normally, granule cells regulate the excitability of DG network with their instinct low firing ability[10, 11]. However, the epileptic granule cells seem to be functional reprogrammed, facilitating the onset of seizure activities[7, 12]. A pioneer study conducted in human hippocampus surgically removed during the treatment of temporal lobe epilepsy showed a functional diversity in DG: the majority of granule cells are hyperexcitable while a small group keeps the normal firing patterns [9]. More recent documents further demonstrated additional epileptic-related changes of intrinsic electrophysiological properties in granule neurons, including the increased leak conductance, less negative resting membrane potential and wider spike waveforms[7]. As Evan A. Thomas et al. indicated in their “genetic sensitivity” analysis simulation, even relatively modest functional changes of granule cells are sufficient to induce seizure activity[13]. Unfortunately, the exact roles of those seizure-induced alternations in TLE are still poorly understood. More systematic studies combined with clinical data, animal models and computational simulations are needed urgently.

The neuronal electrical properties and excitability are controlled by an orchestra of multiple ion channels and metabotropic receptors. Numerous potential epileptic genes code ion channels regulating neuronal firing patterns, although some of them can only explain a small population of idiopathic epilepsies. Among those genes, KCNQ2/3 have been highlighted as having profound connections to the onset of benign familial neonatal seizures in previous documents [14, 15]. At the membrane of granule cells, the protein products coded by KCNQ2 and KCNQ3 assemble heteromultimers, which mediate M-current, a voltage-gated noninactivating potassium current. Activation of M-type channels increases the threshold of action potential, leading to the down-regulation of firing rates during a spike train, termed as spike frequency adaptation (SFA) [16, 17]. A potent SFA is supposed to reduce the fire rate of excitatory neuron during long-term activity. Meanwhile, seizure always couples with the synchronization of neural activities in an epileptic network. Therefore, in epileptic dentate gyrus, SFA of granule cells may prevent the synchronization by decreasing the synaptic positive feedbacks conducted by back sprouting of mossy fibers[5, 11]. However, more compelling investigations are still missing. Combined with the decreased SFA demonstrated in granule cells from TLE patients, it would not be surprising that dysfunctions of KCNQ2/3 may play an important role in the onset of TLE.

In current study, we started from the quantitative analysis of two public databases of TLE patients. Then the correlation among seizure onset and reduced SFA as well as changed KCNQ2/3 expressions were further investigated in the pilocarpine mouse model of TLE and computational simulation of epileptic network, which will provide new insights to understand the mechanisms of epileptogenesis in TLE patients.

## METHODS

### Analysis of human datasets

Two public human TLE datasets were analyzed. The expression data of first one (Database 1) published by Mar Matarin et al.[18] were collected from Gene Expression Omnibus (GEO) with the accession code GSE46706 as well as the authors’ website (http://www.seizubraineac.org/). The second database (Database 2) is from Lars J. Jensen et al.[19] Those data were downloaded from GEO with the accession number GSE134697. All analyses were conducted with RStudio (Version 1.2.1335).

### Pilocarpine-induced mouse model of TLE

Mice were treated with pilocarpine-HCl (Aladdin P129614) via intraperitoneal injection (i.p.) to induce the seizure onset as previously described[20]. For each animal, a single dose of pilocarpine (280–290 mg/kg) in 0.1 ml sterile saline vehicle (0.9%NaCl) was applied. To reduce the peripheral cholinergic effects, 1mg/kg of raceanisodamine hydrochloride (JIANGSU DAHONGYING-HENGSHUN PHARMACEUTICCAL CO, LTD, H32026700) was injected (i.p.) 30 minutes before pilocarpine treatment. Age-matched mice were injected with a comparable volume of vehicle as control. 2-hour behavioral observation after pilocarpine injection was conducted to evaluate the induced seizures, based on a modified version of the Racine’s scale using categories 1–5. Only the mice that experienced at least three category 3–5 seizure onsets were used for further analyses.

### Immunoblotting

Mouse dentate gyrus was dissected as previously described. The tissues were homogenized and lysed in SDS loading buffer, and after quantification, bromophenol blue was added to a final concentration of 0.1%. 25~30ug of total protein was resolved in 10% Nupage Bis-Tris gel and probed with followed primary antibodies: mouse monoclonal anti-KCNQ2 antibody (Proteintech, 66774-1-Ig, 1: 1,000), rabbit polyclonal anti-KCNQ3 antibody (Proteintech, 19966-1-AP, 1: 1,000) and mouse monoclonal anti-beta-actin antibody (Sigma, A5441, 1: 10,000).

### Electrophysiology

Slices including hippocampal dentate gyrus (300 μm) were prepared from 5-week C57/BL6 mice. ACSF (artificial cerebrospinal fluid: 124 mM NaCl, 2.5 mM KCl, 1.2 mM NaH_2_PO_4_, 24 mM NaHCO_3_, 2 mM CaCl_2_, 2 mM MgSO_4_, 12.5 mM Glucose, and 55 mM HEPES; pH 7.35) was prepared before slicing. After anesthesia with chloral hydrate (300 mg/kg) and heart perfusion with cold ACSF, the mice were decapitated with a guillotine. The cortical slices were cut with a vibratome in oxygenized (95% O_2_/5% CO_2_) in ACSF (Sinopharm Chemical Reagent Co., Ltd., Beijing, China) at 4°C and then recovered in NMDG solution (93mM NMDG, 93 mM HCl, 2.5 mM KCl, 1.2 mM NaH_2_PO_4_, 30 mM NaHCO_3_, 20 mM HEPES, 25 mM Glucose, 10 mM MgSO_4_, 0.5 mM CaCl_2_, oxygenized by 95% O_2_/5% CO_2_) at 35°C for 7 minutes. At last, the slices were maintained in normal oxygenated ACSF at 35°C for 0.5 hours and room temperature for 0.5 hours before the experiments. A slice was transferred to a submersion chamber (Warner RC-26G, Molecular Devices, CA, USA) that was perfused with normal ACSF for electrophysiological experiments[21, 22].

We exerted whole cell recording (MultiClamp-700B, Molecular Devices, CA, USA) on dentate gyrus granule cells under a fluorescent/DIC microscope (Nikon FN-E600). Transient capacitance was compensated, and the output bandwidth was 3kHz. The pipette solution contained 150mM K-gluconate, 4mM NaCl, 0.5mM ethylenebis (oxyethylenenitrilo) tetraacetic acid, 4mM Mg-ATP, 1mM Tris-GTP, 5mM Na-phosphocreatine, and 10mM hydroxyethyl piperazine ethanesulfonic acid (adjusted to pH 7.4 by 2M KOH) (Sinopharm Chemical Reagent Co., Ltd.). The osmolarity of the freshly made pipette solution was 295–305 mOsmol, and the pipette resistance was 10–15 MΩ.

### Computational simulation

A simplify dentate gyrus network was simulated by NEURON (v7.2), which only included two types of neurons: granule neuron (GN) and inhibitory neuron (IN). Both of them were adopted from previous published models[5, 11, 23]. In our simulation, granule neuron was composed by a single compartment and only three types of voltage-dependent currents were considered, including fast sodium and potassium currents for the generation of action potentials and M-type potassium current (kinetics from previous reports[24, 25]) for spike frequency adaptation. Other ion channels such as voltage-dependent calcium currents and calcium dependent potassium currents were ignored in this model. For inhibitory neurons, we used the fast-spiking interneuron as representation. Each inhibitory neuron was also composed by a single compartment. But not like the granule neuron, only fast sodium and potassium currents were taken into consideration. Since it has no M-type current, the inhibitory neuron didn’t show spike frequency adaptation. The details of passive and channel parameters were listed in the following table.

In terms of M-type current, it was described by following equation:

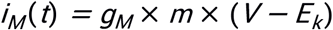

Where

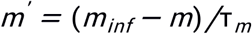

In this equation, minf and τm are both the functions of membrane potential (V):

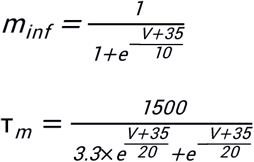

The neurons in our simulated network were connected by chemical synapses. The excitatory synaptic currents were conducted by two types of receptors: a-amino-3-hydroxy-5-methyl-4-isoxazolepropionic acid (AMPA) receptor and N-methyl-D-aspartic acid (NMDA) receptor, while only fast GABAergic synapse (GABA_A_R) was taken into consideration as inhibitory synapse. The excitatory post-synaptic conductance and inhibitory post-synaptic conductance were both described by a double-exponential function [26, 27],

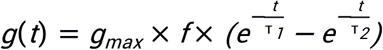

Here f is scale factor, which was calculated by:

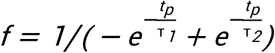

And

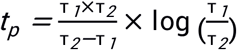

Values of parameters in these equations were listed in table 2.

**Table 2.** passive and channel parameters

| | $Ra$<br>($\Omega cm$) | $C_m$<br>($pF/cm^2$) | $E_{pas}$<br>($mV$) | $g_{pas}$<br>($mS/cm^2$) | $E_{Na}$<br>($mV$) | $g_{Na}$<br>($mS/cm^2$) | $E_K$ ($mV$) | $g_K$<br>($mS/cm^2$) |
| --- | --- | --- | --- | --- | --- | --- | --- | --- |
| GN | 100 | 1 | -70 | 0.1 | 50 | 50 | -100 | 5 |
| IN |  |  |  | 0.15 |  |  |  | 10 |

**Table 2.** Parameters of synaptic connection

| | $\tau_1 (ms)$ | $\tau_2 (ms)$ | $E (mV)$ | $g_{max} (nS)$ | | | |
| --- | --- | --- | --- | --- | --- | --- | --- |
|  |  |  |  | PF→GN | GN→GN | GN→IN | IN→GN |
| AMPA | 1 | 2 | 0 | 0.05 | 0.025 | 0.01 | N/A |
| NMDA | 0.67 | 8 | 0 | 0.001 | 0.005 | 0.005 | N/A |
| GABA<sub>A</sub> | 1 | 6.5 | -75 | N/A | N/A | N/A | 0.02 |

Based on previous study, our network was arranged in a simple linear structure including 96 granule neurons and 32 inhibitory neurons. It meant that in every four neurons there was one inhibitory one. The excitatory neuron received excitatory synaptic input both from the nearby GN and external projection fibers (PF), while was inhibited by neighboring inhibitory neurons. The input from PF followed gamma distribution, which has been well demonstrated by J. Luthman et al.[28] The inhibitory neuron only received the excitatory synaptic input from nearby GN (Fig. 1A). Each excitatory neuron was connected with 4 other neighboring excitatory neurons, and each inhibitory neuron was connected with 6 excitatory neurons nearby (Fig.1B). The weights of these synapses were also listed in table 2.

**Figure 1.**
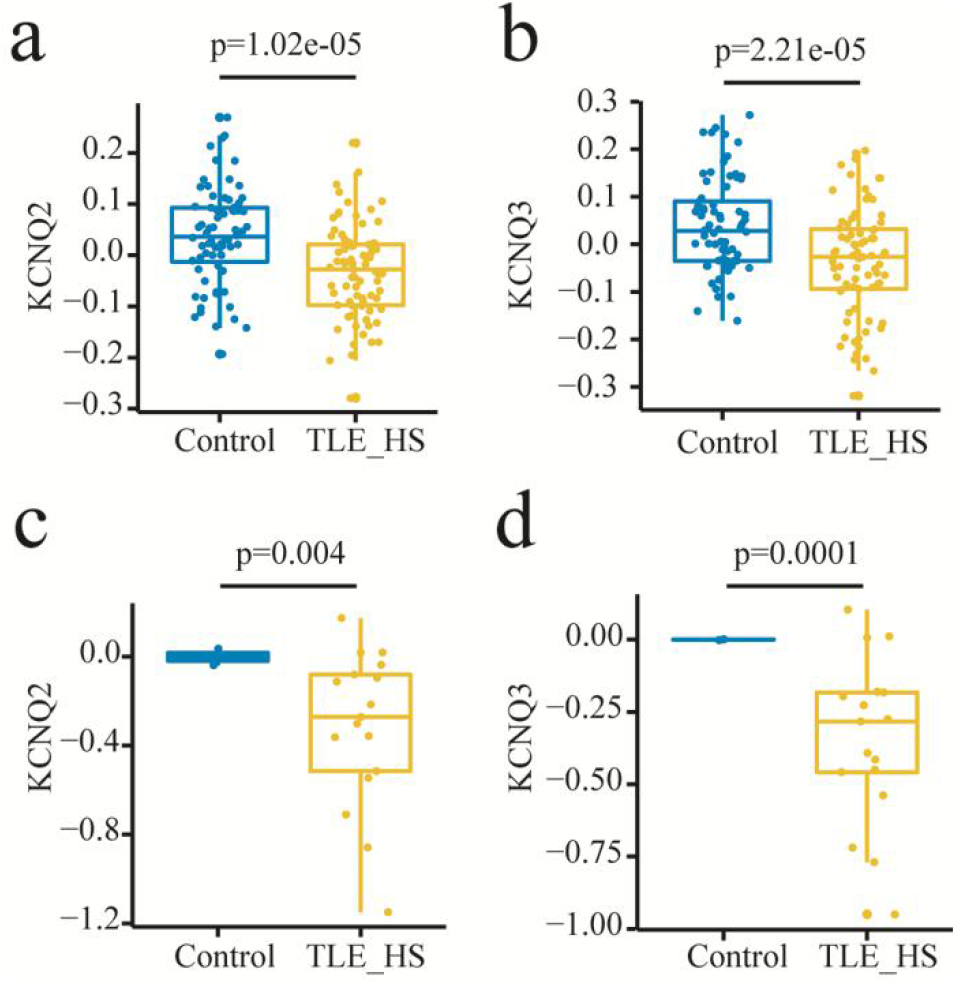
Down-regulation of KCNQ2 and KCNQ3 in TLE patients’ hippocampus. Two gene expression profiling databases both indicate significant decreases of KCNQ2 and KCNQ3 expressions in TLE patients’ hippocampus. **a**, mRNA levels of KCNQ2 in TLE hippocampus vs control, from Database 1 (see Method). **b**, mRNA levels of KCNQ3 in TLE hippocampus vs control from Database 1. **c**, mRNA levels of KCNQ2 from Database 2. **d**, mRNA levels of KCNQ3 from Database 2 (see Method). Significant differences are based on unpaired Students’ t-test. TLE_HS: temporal lobe epilepsy with hippocampal sclerosis.

All the simulations were completed in NEURON[29], the dt was 0.025ms and temperature was 36°C

## RESULTS

### Down-regulation of KCNQ2/3 in hippocampus of TLE patients

Encouraged by progress of high-throughput screening technologies, the searching of epilepsy-associated genes has gained significant momentum over last few years[30–32]. However, most of studies depend on the large genome-wide association study, which is insufficient to address the correlation between alternations of gene expressions and the onset of epilepsy. Several databases of global expression profile derived from TLE patients are published recently, providing feasible resources to identify novel candidates of epileptic genes. We first examined the expression changes of KCNQ2/3 in newly published transcriptomic dataset recruiting 85 TLE patients with hippocampal sclerosis and 75 healthy controls with no neurological or neuropsychiatric disorders[18]. Both mRNAs of KCNQ2 and KCNQ3 showed significant decreases in TLE patients (Fig. 1a and b). We also looked into the variation of different transcripts and only found a significant change of KCNQ2 but not KCNQ3, indicating a potential impact of altered splicing patterns (data were not shown). Next we confirmed this correlation in another dataset including 17 TLE patients as well as 2 healthy controls, which was published by Lars J. Jensen et al[19]. Although the sample size is relatively small, the decreases of KCNQ2/3 are still statistically significant, as shown in Fig. 1c and d. Interestingly, we found weak positive correlations between KCNQ2/3 and duration of epilepsy (Fig. 2a) from the second database, indicating a possible adaptive plasticity coupling with the progress of epilepsy, which is consistent with previous reports.[8] Meanwhile, the weak downhill relationships between KCNQ2/3 levels and onset age but not current age (Fig. 2b and c) may suggest the gradually loss of abilities to homeostatically rescale intrinsic physiological properties in response to the onset of seizure.

**Figure 2.**
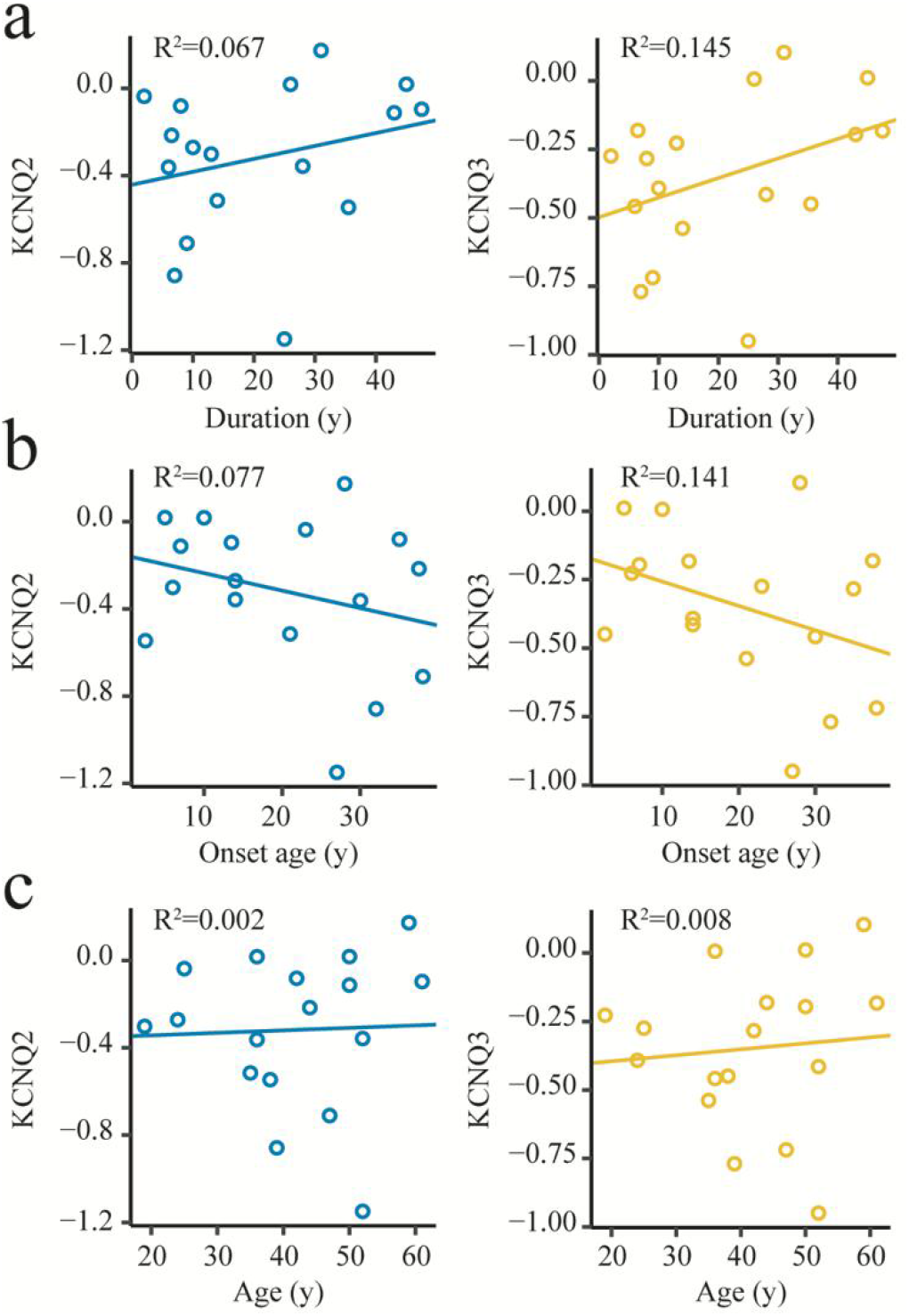
Correlations between KCNQ2/3 expression and patients’ characteristics. The relationships of mRNA levels of KCNQ2/3 and duration of epilepsy (**a**), onset age (**b**) as well as age (**c**) in Database 2 were exhibited respectively.

### Seizure induced dysfunction of dentate granule cells

To further investigate the functional outcome of down-regulation of KCNQ2/3 evoked by seizure, we employed a well-characterized chemical-induced TLE mouse model. The mice were treated with pilocarpine (280–290 mg/kg, i.p.) and evaluated by Racine’s scale[20]. Animals with seizure activities that met the criteria described before were euthanized for analysis of protein expressions and electrophysiological properties. The significant decreases of protein levels of KCNQ2 and KCNQ3 in the epileptic dentate gyrus were determined by immunoblotting (Fig. 3a and b), which is in line with the patterns we demonstrated in human datasets. The similar decreases of KCNQ2/3 proteins in hippocampus from both patients and TLE mouse model or mice indicates that the pilocarpine-induced model faithfully represents the pathophysiological process associated with M-currents potassium channels. Wealthy literatures have documented the crucial roles of KCNQ2/3 in the regulation of neuronal excitability[33, 34]. Therefore, we continually examined the potential electrophysiological dysfunctions of granule cells affected by reduced KCNQ2/3. Slices with hippocampal DG area were prepared to conduct the recording of granule cells. Granule cells were selected according to the localization and morphological properties, which were further confirmed by the staining of the fluorescent dye Alexa 488 loaded in the pipette solution. As expected, the same stimulation induced more spikes in the epileptic granule neurons in comparison with healthy control cells (Fig. 3c). We further analyzed the initial frequency (f_0_) calculated as the reciprocal of the first inter-spike intervals (Fig. 3d), as well as the steady-state frequency (f_inf_, as shown in Fig. 3e). Only a slight increase of initial spike frequency was observed in epileptic granule neurons, but the difference in these 2 groups failed to reach the statistical significance (p=0.07, Fig. 3d). In contrast, the gain of steady-state frequency in epileptic group was significant (Fig. 3e), suggesting the down-regulation of spike frequency adaptation. Altogether, we determined the reduced expression of KCNQ2/3 and impacted SFA in mouse dentate granule neurons after epileptic seizure induced by pilocarpine, further confirmed the correlation of TLE onset and the dysfunction of M-current potassium channels identified in patients’ hippocampus.

**Figure 3.**
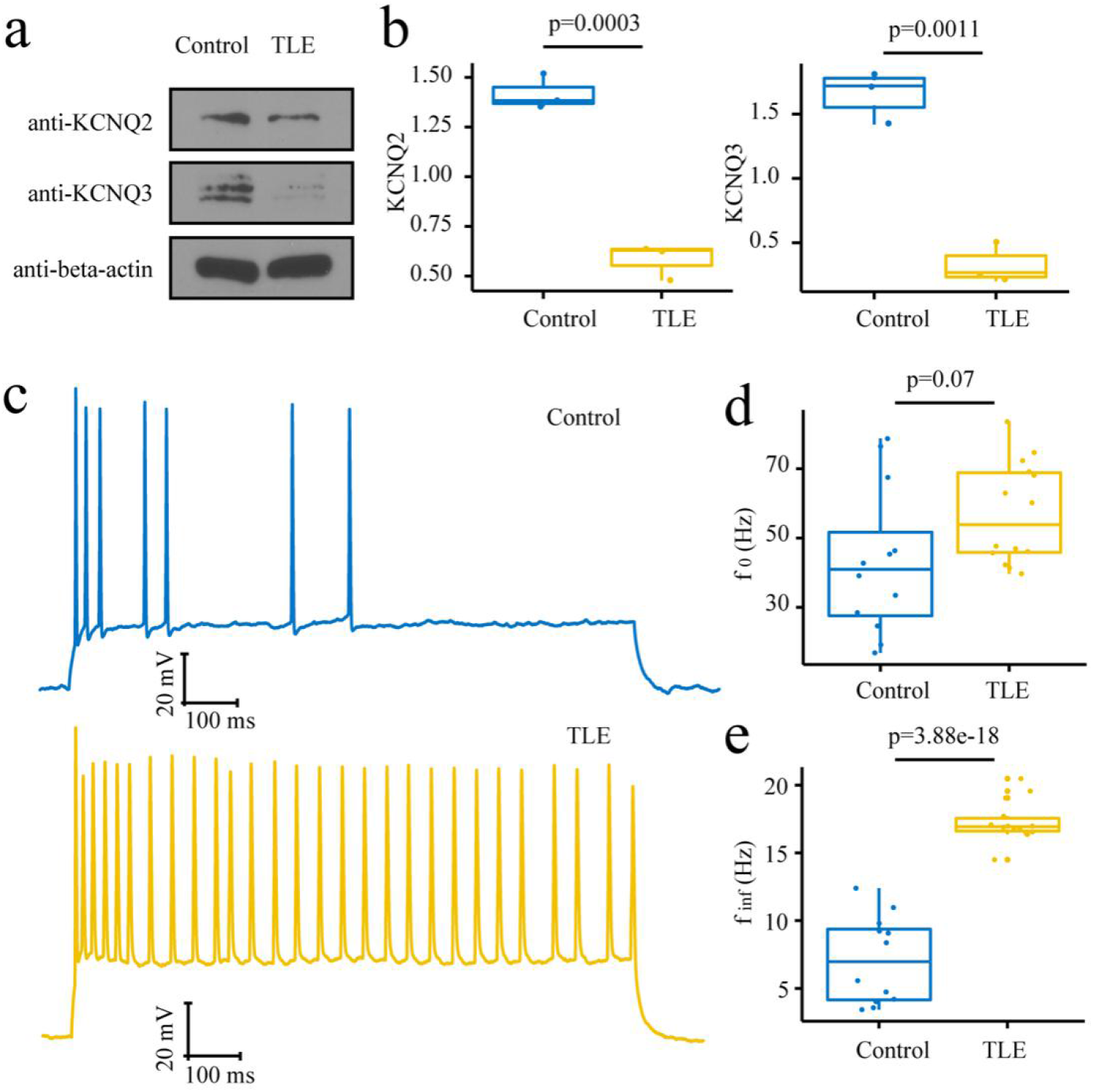
Impacted KCNQ2/3 expressions and SFA in pilocarpine-induced TLE mouse model. **a**, decreased protein levels of KCNQ2/3 in dentate gyrus of TLE mouse were detected by immunoblotting. **b**, statistical analysis of **a**. **c**, representative traces of spike firing captured from granule cells in control (blue) or TLE dentate gyrus. **d** and **e**, statistical analysis of initial frequency (f_0_, in panel **d**) and steady-state frequency (f_inf_, in panel **e**). All significant differences are based on unpaired Students’ t-test.

### Reduction of spike frequency adaptation facilitates seizure activities

Previous studies have suggested the potent spike frequency adaptation of dentate granule neurons is crucial in filtering the input signals towards hippocampus[11]. The low firing ability of granule cells regulates the excitability of entire network and consequently, may prevent possible abnormal hyperactivities. However, the exact effect of M-currents conducted by KCNQ2/3 in an epileptic hippocampus network has not been completely demonstrated. Therefore, we employed a computational simulation to perform the following studies that currently are extremely difficult using other approaches.

To stimulate the distinct fire patterns of granule cells under epileptic or control condition, we firstly validated the relationship between M-currents and SFA in our model neurons. Without M-currents (g_M_=0), the firing frequency of model neuron was fixed, while a significant adaptation appeared coupling with the increase of conductance of M-currents channels (Fig. 4a). The instantaneous frequency defined as the reciprocal of inter-spike interval was plotted in Fig. 4b, showing a typical SFA regulated by M-currents. To compare with the spike firing patterns which we documented in TLE mouse model, the initial frequency (f_0_) and steady-state frequency (f_inf_) were exhibited in Fig. 4c and d respectively. The decreases of f_0_ and f_inf_ induced by the repression of M-currents is comparable to the electrophysiological dysfunctions of epileptic granule cells, confirming that our model cell reproduced the key pathological alternations associated with the seizure-induced reduction of KCNQ2/3.

**Figure 4.**
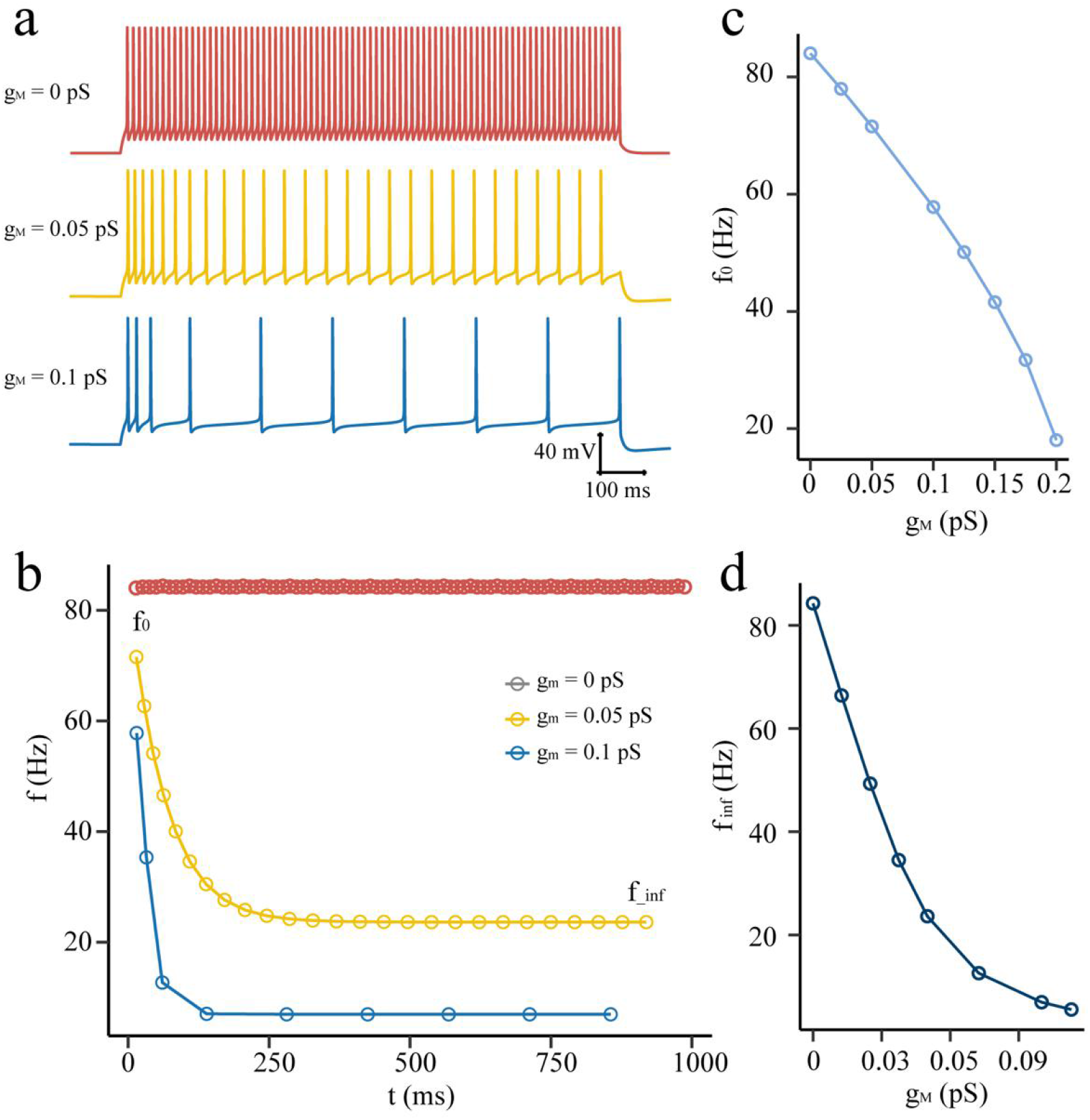
Spike frequency adaptation of model granule neuron is regulated by conductance of M-type channels. **a**, the spike fire patterns of model granule neuron with different density of M-channel conductances. **b**, the instantaneous frequencies of model neurons gradually decrease from the initial frequency (f_0_) down to the steadystate frequency (f_inf_) upon the activity of M-type channels. **c** and **d**, the initial frequency (**c**) and steady-state frequency (**d**) exhibit negative correlations with conductance of M-channnels.

Next we composed a simplified dentate gyrus network only containing granule cells and inhibitory neurons. The sprouting mossy fiber was represented by the excitatory synaptic connections between granule cells, which is not exist in non-epileptic hippocampus (see Method). Unprovoked epileptiform activities, characterized by high spike frequency and synchronization between granule cells, burst repeatedly in our simulated network even when the conductance of M-currents is not impacted (g_M_=0.11 pS/cm^2^, Fig. 5a). Such hyperexcitability of dentate network is the consequence of enhanced positive feedbacks between granule cells conducted by mossy fiber sprouting, which is in the same line with previous studies[4–6]. However, the seizure would not last forever and an interictal pause appeared between two seizures. In this period, granule cells still generated sporadic action potentials, but would not evoke the synchronized activity of whole network. Thus, our simulated dentate gyrus represented the key features of epileptic network. Next, we found more seizure onsets were appeared coupling with the repression of M-currents (g_M_=0.05 pS/cm^2^, Fig. 5a). If the adaptation of frequency was almost entirely abolished, the epileptiform activities appeared very soon after the beginning of simulation and continued without a stop (g_M_=0.01 pS/cm^2^, Fig. 5a). The right panel of Fig. 5a displays the firing pattern of a representative model of granule cell, showing high frequency burst of action potential that was synchronized with the seizure spreading over the whole network.

**Figure 5.**
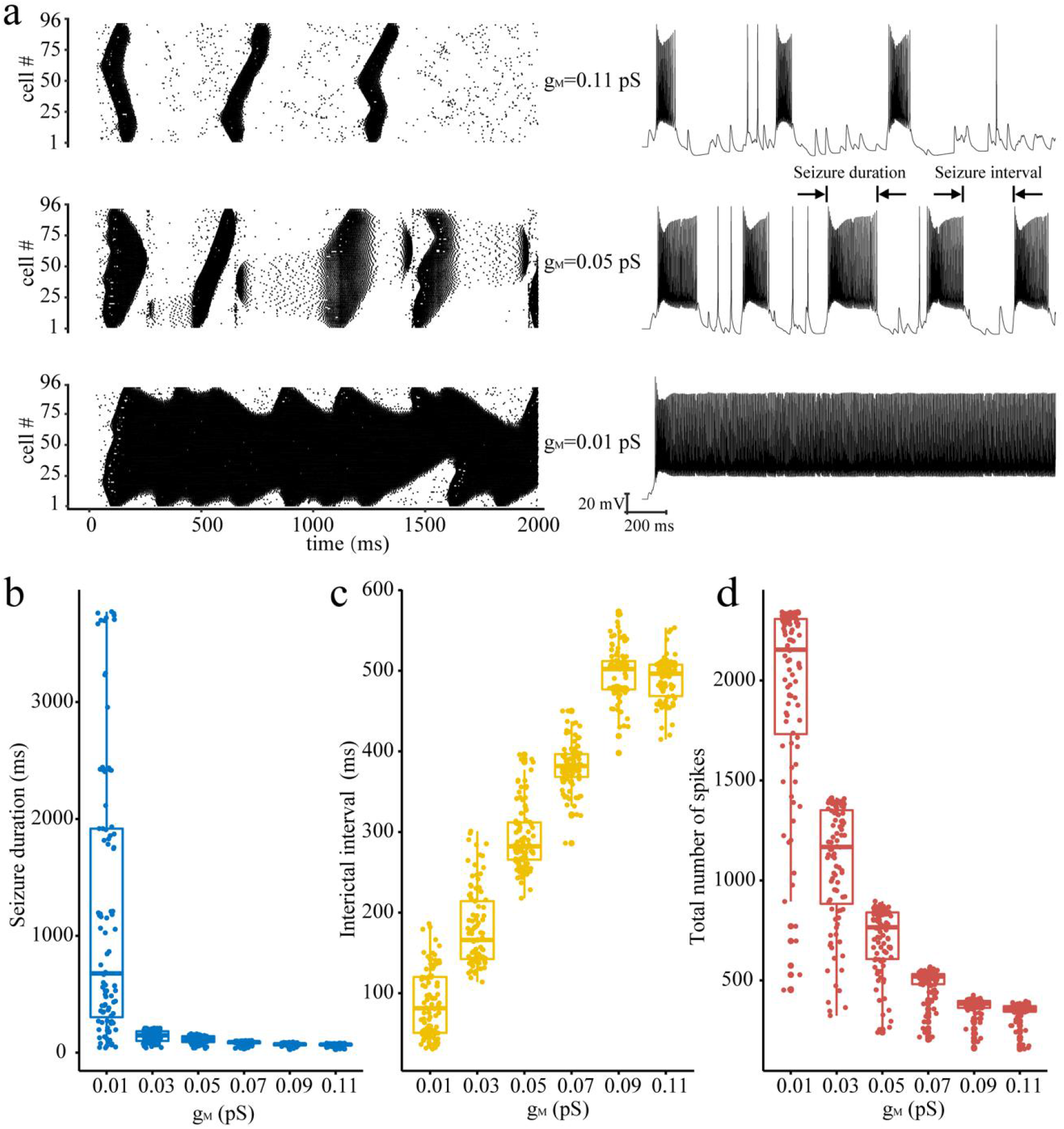
Decreased M-currents facilitated the onset of epileptiform activities in simulated dentate network with mossy fiber back sprouting. **a**, Action potential firing of model granule cells coupling with different density of M-type channels. In the left panels, each dot indicates a spike event. The right panel shows the membrane potential of one neuron as representative trace. **b** to **d**, the density of M-type channels controls the properties of seizure activities in simulated dentate network, including seizure duration (**b**), interictal interval (**c**) and total number of spikes (**d**).

To evaluate the correlation between SFA of granule cells and seizure activities of simulated dentate network, plots of g_M_ versus seizure duration, interictal interval and total spike number were presented in Fig. 5b, c and d respectively. The increase of spike frequency adaptation in granule cells prolonged interictal durations and shortened seizure durations. These results demonstrate that the SFA of excitatory neurons in network helps to prevent the long-term persistence of seizure activity. However, the spike frequency adaptation cannot prevent seizure in the range of g_M_ set in our simulation. A higher g_M_ (>0.3pS/cm2) would fully eliminate the spontaneous seizure onset in this network (data were not shown).

In conclusion, employing pilocarpine-induced TLE mouse model and computational simulation we confirmed that the decreased expression of KCNQ2/3 impacts the spike frequency adaptation in epileptic granule cells, thus facilitates the onset of epileptiform activities among the entire dentate gyrus network. Our results represent the first functional analysis of KCNQ2/3 and spike frequency adaptation in epileptic dentate granule cells, suggesting a novel mechanism in TLE dentate gyrus.

## DISCUSSION

The heterotetrameric channels formed by the products of KCNQ2 and KCNQ3 conduct M – currents, regulating the spike fire pattern and excitability of neurons. Mutations of KCNQ2 and KCNQ3 are originally reported as the leading drivers inducing familial seizures in infants, suggesting the essential function of M-currents potassium channels in repressing of network hyperexcitability. However, the transgenic mouse model expressing mutated KCNQ2 or KCNQ3 did not exhibit the typical TLE-related pathology including loss of hippocampal neurons and sprouting of mossy fibers[35]. Only changed intrinsic membrane properties of CA1 pyramidal cells were documented. Those results raise more questions about the possible roles of KCNQ2/3 in TLE hippocampus. In the first part of our study, we focused on two public databases and documented the significant decreases of KCNQ2/3 expressions, providing strong evidence to show the intimate link between those two genes and seizure susceptibility in TLE patients. However, limited by the restriction of human samples, it is still unclear which subtype of neurons in hippocampus are affected by decreases of KCNQ2/3 expressions. Thus by the employment of TLE mouse model and computational simulation, we demonstrated that the seizure-induced down-regulations of KCNQ2/3 expressions in dentate granule cells is one of the leading drivers facilitating the hyperexcitability of hippocampus network. Considering a newly reported pharmaceutical activator of M-type channels (ML213)[36], it’s eminently expected that KCNQ2/3 highlighted by our results would become a potential druggable target for the TLE treatment.

Temporal lobe epilepsy is a chronic illness always lasting for many years, associated with multiple long-term neurochemical disturbances and reorganizations of neuronal circuit. Investigation of the pathological changes underlying provoked seizure onset is crucial for the drug discovery of anti-epileptic therapy. Our study focusing on KCNQ2/3 strengthens the significance of seizure-associated expressional changes besides genetic mutations identified by global screening. There is no doubt that recurrent epileptiform activities would affect numerous genes and stimulate a cascade of expressional rearrangements in dentate gyrus and entire hippocampus. The early studies focused on genes associated with axonal sprouting and reorganization of hippocampus network, for instance, GAP-43 and dynorphin [37]. More epileptic-induced expression changes were captured by multiple global expression profiling assays including microarray, serial analysis of gene expression and RNA-seq[31]. However, the mechanisms underlying those expression shifts, including decreases of KCNQ2/3 expression, are still unknown. Therefore, the resetting of epigenetic landscape, which takes part in the chronic pathological process that reforming the epileptic granule neurons and network, is a promising direction for the future investigations. More interestingly, although most samples from TLE patients showed decreased KCNQ2/3 expressions, a weak positive correlation between KCNQ2/3 mRNA and epilepsy duration was detected. It may indicate an endogenous mechanism enhancing the expression of M-currents potassium channels, which rescales the intrinsic membrane property to prevent further seizure onset. A similar plasticity of hyperpolarization-activated current (Ih), was also documented by J. Wolfart et al[12]. The promoted Ih-currents enhance membrane conductance and consequently increase the energy needed for action potential firing. Inspired by those results, the potential therapeutic strategies against TLE would not only limit to revise the down-regulations of KCNQ2/3 expressions but also include approaches promoting the homeostatical plasticity of hippocampus neurons.

## Acknowledgments

This study is funded by grants from the Natural Science Foundation China (31500836), Natural Science Foundation of Anhui Province (1608085QH176) and the Key Program of Anhui Educational Committee (KJ2019A0299) to Rongjing Ge.

## Authors’ contributions

R. G. and H. Q. designed the whole project. S. J., R. L., S. T., and B. L. did the experiments and data analysis. B. L. and H. Q. worked on computational simulation. R. G., H. Q., and F. M. wrote the manuscript. All authors have read and approved the final manuscript.

